# An explainable graph neural framework to identify cancer-associated intratumoral microbial communities

**DOI:** 10.1101/2023.04.16.537088

**Authors:** Zhaoqian Liu, Yuhan Sun, Anjun Ma, Xiaoying Wang, Dong Xu, Daniel Spakowics, Qin Ma, Bingqiang Liu

## Abstract

Microbes are extensively present among various cancer tissues and play a vital role in cancer prevention and treatment responses. However, the underlying relationships between intratumoral microbes and tumors are still not well understood. Here, we developed a MIcrobial Cancer-association Analysis using a Heterogeneous graph transformer (MICAH) to identify intratumoral cancer-associated microbial communities. MICAH integrates metabolic and phylogenetic relationships among microbes into a heterogeneous graph representation. It uses a graph attention transformer to holistically capture the relationships between intratumoral microbes and cancer tissues, which improves the explainability of the association between identified microbial communities and cancer. We applied MICAH to intratumoral microbiome data across five cancer types and demonstrated its good generalizability and reproducibility. We believe this graph neural network framework can provide novel insights into cancer pathogenesis associated with the intratumoral microbiome.

## INTRODUCTION

Microbes have been detected as an intrinsic component of the tumor microenvironment across multiple cancer types^1-3^. These commensal microbes closely interact with each other to perform functions synergistically, such as producing metabolites that promote tumor growth or stimulating the immune system to attack cancer cells^4,5^. Compelling evidence indicates that each cancer type has a unique microbiome, characterized by microbial communities with specific functions^6^. These microbial communities are highly associated with cancer initiation, progression, and pathogenesis^2,3,6-9^. A comprehensive characterization of the cancer-associated microbial communities facilitates a mechanistic understanding of the links between intratumoral microbiome and cancer. Moreover, it will shed light on translating microbiome research to clinical diagnosis and treatment via manipulating intratumoral microbiome^8^. However, the exploration of the links between intratumoral microbiome and cancer tissues is far from complete due to the difficulties in obtaining clinical biopsies specifically dedicated to microbial profiling^10^. Fortunately, the advances in sequencing technologies provide large-scale human tissue sequencing data, such as The Cancer Genome Atlas (TCGA) and the Oncology Research Information Exchange Network^11^. Several computational tools, such as PathSeq and SRSA^12,13^, have been proposed to mine intratumoral microbial reads from existing tissue sequencing data, which enables the characterization of the tissue-resident metagenome without the need for additional experimental manipulation^1,3,10,14,15^. The resulting data, such as The Pan-Cancer Mycobiome Atlas and The Cancer Microbiome Atlas (TCMA)^1,10^, serve as valuable resources for exploration of the intratumoral microbiome.

Existing approaches for identifying disease-associated gut microbiomes can be. transferred to the study of cancer-associated intratumoral microbiomes^3,16,17^ These approaches generally identify microbial dysbiosis between normal and tumor tissues as disease-associated microbial signatures through comparative studies^6^. However, the number of precisely matched solid tissue normal samples of cancers is generally limited, especially in TCGA, which greatly limits the application power of such studies. Another concern is that comparative studies between normal and tumor tissues do not contextualize other diseases, potentially resulting in identified microbiomes that are not specific to a particular disease but rather reflect shared patterns across multiple diseases or common symptoms^18^. This hinders the mechanistic understanding of a specific disease and future research from association to causation^18^. From a methodological perspective, there are still several issues with the existing approaches. The most widely used method is to identify key microbes as signatures based on differential abundance of individual microbes between two groups of cohorts^14,19-22^, or by measuring the relevance between the abundance of individual microbes and sample labels using traditional machine learning^23^. However, these methods ignore the intricate interactions that reflect the structure and functions within microbial communities^24,25^. Although several studies have attempted to address this issue by constructing co-occurrence networks^24,26^, obtaining a reliable network that can represent the true species-species interactions is still a challenge^27^, which is prone to bias in downstream analysis. These issues have motivated us to design effective methods to discover high-confidence cancer-associated microbial communities, thereby enhancing mechanistic insights into the relationships between the microbiome and cancer.

In this study, we proposed a MIcrobial Cancer-association Analysis using a Heterogeneous graph transformer (MICAH) to identify intratumoral microbial communities associated with cancer. By integrating complex relationships among microbes and between microbes and hosts, our method can provide insights into elucidating the biological mechanisms underlying cancer-microbiome relationships. Specifically, we constructed a heterogeneous graph to represent the intricate relationships between microbes and their hosts, where the phylogenetic and metabolic relationships among microbial species were also included as knowledge edges in the initial representation. Based on this heterogeneous graph representation, we deployed a graph attention transformer to capture the links between microbes and multiple cancer types, and holistically interpreted the relevance between microbial communities and sample phenotypes. Finally, we identified cancer-associated microbial communities for each cancer type by selecting communities consisting of statistically significant species with high attention scores. We applied MICAH to a dataset of five cancer types in TCMA^10^ (Colon adenocarcinoma, abbr. COAD; Esophageal carcinoma, abbr. ESCA; Head and Neck squamous cell carcinoma, abbr. HNSC; Rectum adenocarcinoma, abbr. READ; and Stomach adenocarcinoma, abbr. STAD), demonstrating its high accuracy and generalizability in cancer classification, as well as high reproducibility in identification of microbial communities associated with cancer. We believe the proposed graph neural network (GNN) framework is highly effective for identifying cancer-associated microbial communities and has the potential to impact our understanding of cancer pathogenesis, diagnosis, and treatment.

## METHODS

### Datasets

We retrieved intratumoral microbial profiles from the TCMA database, which provided microbial abundance data for 512 samples from primary tumor tissues across five TCGA projects (COAD: 125 primary tumors, ESCA: 60 primary tumors, HNSC: 155 primary tumors, READ: 45 primary tumors, and STAD: 127 primary tumors) ^10^. We extracted microbial profiling at the species-level resolution and generated a species-sample abundance matrix comprising 1,218 microbial species from the 512 samples.

### MICAH: A framework to identify cancer-associated intratumoral microbial communities

MICAH is an end-to-end GNN framework for cancer-associated microbial community inference. This framework leverages a heterogeneous graph transformer (HGT) model to classify samples and employs attention scores obtained from the model to evaluate the relevance between microbial species and cancer types. MICAH takes known cancer labels for samples, represented as a vector *Y*, and a microbial abundance matrix, *A*, as required input for the model training, and metabolic and phylogenetic relation matrices as optional input. The abundance matrix is organized with species as rows and samples as columns, where each element denotes the abundance of a particular microbial species within its corresponding sample. The metabolic/phylogenetic relation matrix indicates whether there is a metabolic/phylogenetic relationship between two species, with a value of one indicating a relationship and zero indicating no relationship. The output of MICAH is the microbial community of each cancer type, which can be prioritized as potential candidates associated with cancer occurrence, development, and treatment for future mechanistic studies.

#### Step 1: Identify nodes and edges from the input abundance matrix

For the input abundance matrix, we conduct a pre-processing first: the rows that contain less than 0.05% non-zero values are removed. Then, we re-normalized the matrix and obtained a new relative abundance matrix, *A*_*M*×*N*_. The abundance matrix used later refers to this preprocessed relative abundance matrix unless exceptions are mentioned.

We used microbial species and cancer samples within the abundance matrix as species and sample nodes, respectively. The species-sample edges are constructed based on the present abundance of species within the sample. If the relative abundance is non-zero, there will be an edge between the species and the sample. The species-species edges are constructed based on metabolic and phylogenetic relationships among microbial species. Since microbial metabolomics data of most samples are not available^28^, we rely on an existing database NJS16^29^, which includes more than 4,400 known metabolic interactions between 570 well-studied microbial species, to assess metabolic relationships among species. The NJS16 database collected the experimentally verified metabolic relationships from the literature and listed the compounds produced or consumed by each microbial species. We estimate metabolic relationships between species based on their associated compounds^30^: If two species consume the same compound, there is metabolic competition between the two species^30^; if a compound that one species needs to consume is exactly the one produced by the other species, there is a metabolic complementary relationship between the two species^30^. No matter the metabolic competition or complementarity between two microbial species, we set a metabolic edge between the two species. In this way, we can obtain a metabolic relation matrix, 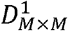, to represent metabolic edges between species, where 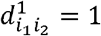 if there is a metabolic edge between the *i*_1_^*th*^species and the, *i*_2_^*th*^ species (*i*_1_ ≠ *i*_2_); otherwise, 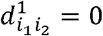.

Phylogenetic relationships are measured using the NCBI taxonomy database. This database includes a hierarchical system of classification that is based on the principles of phylogenetic relationships, with each level representing a different degree of relatedness. We consider the phylogenetic relationships that two species are in the same genus^31^. In this way, a phylogenetic relation matrix, 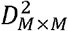, can be obtained, where 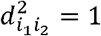 if the *i*_1_^*th*^ species and the *i*_2_^*th*^ species (*i*_1_ ≠ *i*_2_) are in the same genus; otherwise, 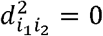.

For the TCMA dataset used in this study, there were 512 sample nodes and 1,218 species nodes, a total of 13,777 metabolic edges and 17,323 phylogenetic edges among these species nodes.

#### Step 2: Construct a microbial species-sample heterogeneous graph

We first introduce the representation of a heterogeneous graph *G* = *(V,E, P,Q)*, where *V* is a node set, *E* is an edge set, *P* and *Q* represent a node type union and an edge type union, respectively^32^. Based on the abundance matrix *A*_*M*×*N*_ and the obtained relation matrices, 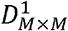 and 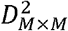, we construct a heterogeneous graph *G* with two node types (microbial species and samples) and three edge types (species-sample edges and species-species metabolic and phylogenetic edges, respectively). Mathematically, *V* = *V*^*s*^ ∪ *V*^*p*^, where 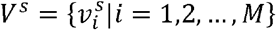 denotes all microbial species nodes, and 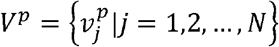 denotes all sample nodes. 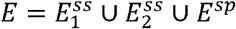, where 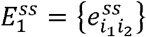 represents metabolic edge unions between two species 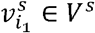 and 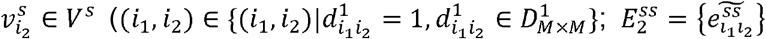 represents phylogenetic edge unions between two species 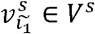 and 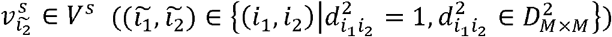; and 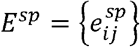 represents the edgesbetween a species node 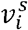 and a sample node 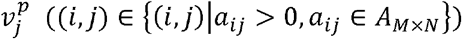.

#### Step 3: Train a heterogeneous graph transformer for cancer sample classification

We propose a supervised GNN model and introduce node- and edge-type dependent attention mechanisms to assess the relevance between microbial species and samples.

Considering the abundance matrix *A*_*M*×*N*_ is extremely sparse and noisy^33^, we first use two autoencoders to extract representative information as the initial embeddings of species and sample nodes in *G*, respectively. The species autoencoder learns low-dimensional features as species nodes embedding, which consists of an encoder with the transpose of *A*_*M*×*N*_, denoted as 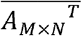, as the input layer and a hidden layer with 256 neurons, and a decoder to restoring the features in the hidden layer. The output layer is a reconstructed matrix 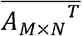 of the same dimension with 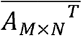. The mean square error between 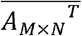 and 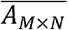 is used as the loss function of the species autoencoder. Similarly, the sample autoencoder learns low-dimensional features as sample nodes embedding, whose structure is the same as the species autoencoder, but the input is the matrix *A*_*M*×*N*_, and the output is 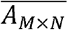 of the same dimension with *A _M×N_*. The loss function is the mean square error between *A*_*M*×*N*_ and 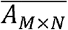. After such processing, we assign a 256-dimensional feature vector to each species and each sample node as the initial embedding.

A GNN model is applied to classify sample cancer types. The input is the adjacency matrix of graph *G* and a feature matrix consisting of node embedding from autoencoders. *L* hidden layers are used to transmit and update the node features, and the default value of *L* is 2 in this work. A fully connected layer with *X* neurons is then connected, in which *X* is the number of cancer types of all samples. The final output is predicted sample labels *Ŷ*. To alleviate the issue of data imbalance, we use Focal Loss as the classification loss function. Given the graph heterogeneity, the node- and edge-type dependent attention mechanism is specifically introduced for node features update in the model^32^. We next focus on how to use attention mechanisms, do message passing and aggregation, and design loss functions.

Before the details, we give several definitions and symbols: 1) the mapping function of node and edge types. We define *τ*(*v*):*V* → *P* and *ϕ*(*e*):*E* →*Q* as the mapping function of node types and edge types, respectively. 2) Source nodes and target nodes. When we focus on a node, the node can be considered as a target node, denoted as *v*_*t*_. If there is an edge between *v*_*s*_ and *v*_*t*_, the node *v*_*s*_ can be considered as a source node of node *v*_*t*_. 3) Meta relation. It describes a connection between a source node *v*_*s*_ and a target node *v*_*t*_, using the node types of *v*_*s*_ and *v*_*t*_, and edge types *e*_*st*_ to represent, denoted as ⟨*τ*(*v*_*s*_), *ϕ*(*e*_*st*_), *τ*(*v*_*t*_)⟩.

##### 3.1 Introducing node- and edge-type dependent multi-head attention mechanism

Attention indicates the importance of a source node *v*_*s*_ to a target node *v*_*t*_. Multi-head attention is a combination of multiple independent attention and helps to attend to parts of the feature differently^34^. We use *h* attention heads and set *h* = 8 as default according to the grid search results.

When we calculate the attention value of a source node *v*_*s*_ to a target node *v*_*t*_, the attention of each head is an independent output, and then concatenated as an attention vector of *v*_*s*_ and *v*_*t*_. Specifically, we denote the embedding of a target node *v*_*t*_ and a source node *v*_*s*_ on the *l*^*th*^ layer as ℋ^*l*^ [*v*_*t*_] and ℋ^*l*^ [*v*_*s*_] (*l* = 0,1, ⋯,*L*), respectively, where ℋ^0^ represents the initial node embedding. *d* is the dimension of ℋ^*l*^ [*v*_*t*_]. For the *k*^*th*^ attention head in the *l*^*th*^ HGT layer, 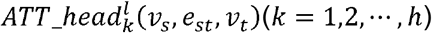 (*k* = 1,2, ⋯*h*), we use node-type-dependent linear projection functions 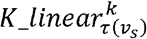 and 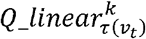 to map the embedding of the source node and target node in the *k*^*th*^ head and (*l* - 1)^th^ layer, respectively, obtaining the *k*^*th*^ Key vector *K*^*k*^(*v*_*s*_) and Query vector *Q*^*k*^(*v*_*t*_) (Fig.1).

**Fig. 1.**
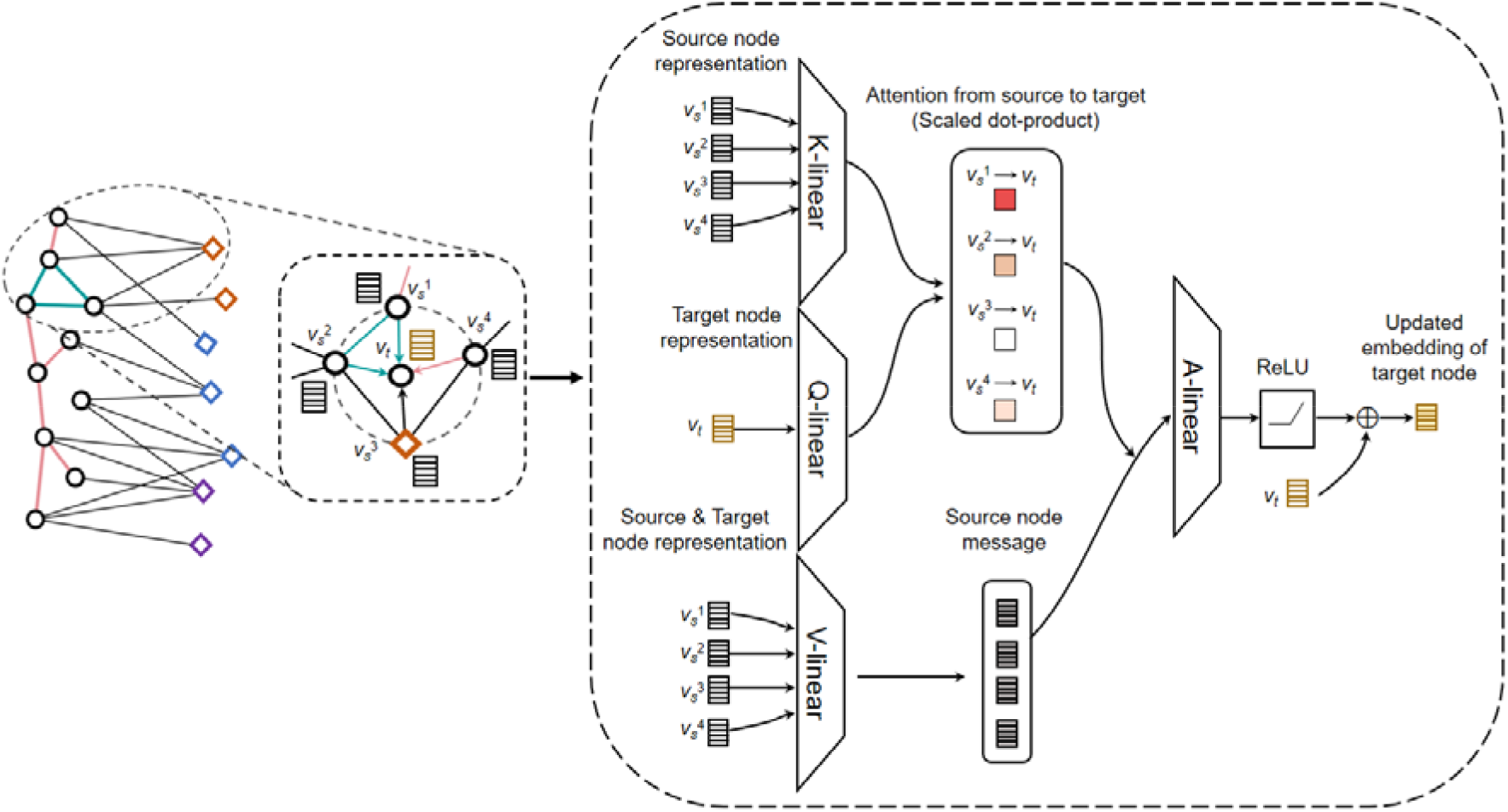
Details of heterogeneous graph transformer. When we update node embeddings, each node is regarded as a target node, and the information of neighbor nodes is used to update the target node embedding. Using a species node *v*_*t*_ as the target node for illustration. Its neighbor nodes, *v*_*s*_^l^, *v*_*s*_^2^, *v*_*s*_^3^ and *v*_*s*_^4^, are considered as source nodes while updating the embedding of the target node. We use node-type-dependent linear projection functions, 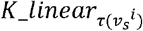(i = 1, 2, 3, 4) and 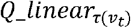, to map the embedding of source nodes and the target node, respectively, obtaining key vectors and a query vector. Here, *τ*(*) indicates the type of a given node. Then, the similarity between each key vector and the query vector is calculated as an attention score from the source node to the target node. Meanwhile, we use node-type-dependent linear projection functions, 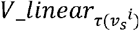, to map the embedding of all source nodes, obtaining value vectors as a message of each source node to the target node. Next, the messages from all source nodes are aggregated and weighted by the corresponding attention scores. By integrating the original node embedding of the target node with the aggregated node messages, we can obtain an updated embedding of the target node.

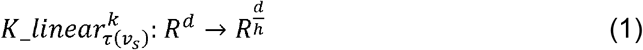

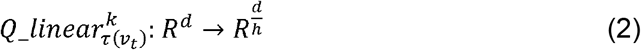

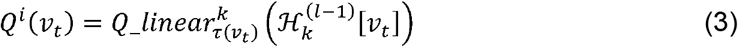

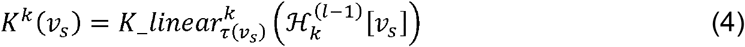

Then, the similarity between the Key vector and Query vector is calculated as 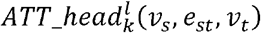. Given the different edge types between the source node and the target node, an edge-type-dependent matrix 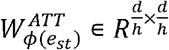 is applied here to capture distinct relationships even between the same node type pair.

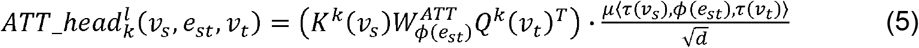

where *μ*⟨*τ*(*v*_*s*_), *ϕ*(*e*_*st*_), *τ*(*v*_*t*_)⟩ is a prior tensor to denote the significance of each meta relation ⟨*τ*(*v*_*s*_), *ϕ*(*e*_*st*_), *τ*(*v*_*t*_)⟩, serving as an adaptive scaling to the attention^32^. Next, the multiple attention heads are concatenated and an attention vector for each node pair is obtained, denoted as 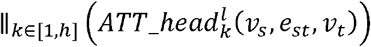, where 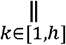 is a concatenation function. Finally, we gather all attention vectors from all source nodes of a target node *v*_*t*_ and use softmax, as Formula (6). In this way, 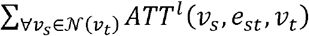 is equal to 1, which facilitates the measurement of the importance of a source node *v*_*s*_ to a target node *v*_*t*_.

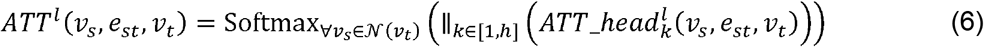

where 𝒩(*v*_*t*_) is a set of source nodes of *v*_*t*_

##### 3.2 Message passing and aggregation

In this step, we first extract the message passed from a source node *v*_*s*_ to a target node *v*_*t*_. Then, we aggregate the message from all source nodes of *v*_*t*_ using the corresponding attention as weight, obtaining updated embedding for the target node *v*_*t*_.

Specifically, for the *k*^*th*^ message head in the *l*^*th*^ HGT layer from *v*_*s*_ to *v*_*t*_, 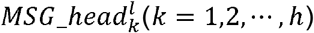, we use a node-type-dependent linear projection function, 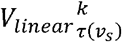, to map the embedding in the *k*^*th*^ head and (*l* – 1)^*th*^ layer, obtaining the *k*^*th*^ Value vector *V*^*k*^(*v*_*s*_) (Fig.1).

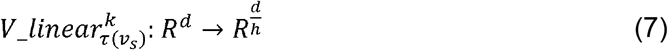

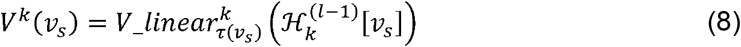

Incorporating an edge-type-dependent matrix 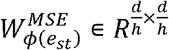, we use the Formula (9) to

Compute 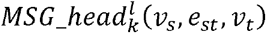. All *h* message heads are concatenated for the overall message of the node pair *v*_*s*_ and *v*_*t*_, *MSG*^*l*^ (*v*_*s*_,*e*_*st*_, *v*_*t*_).

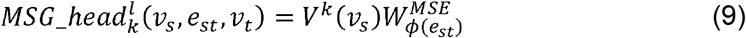

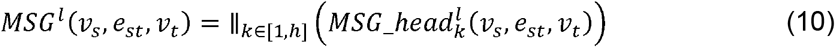

Finally, for a specific target node *v*_*t*_, the message of all of its source nodes is aggregated for embedding update (Formula (12), Fig.1). The attention (after the softmax function) is used as the weight of the passing message from the corresponding source node, *θ* is a trainable parameter, and *ReLU* is the activation function.

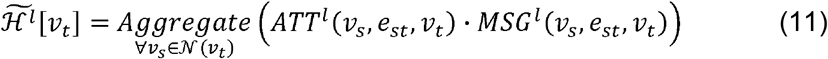

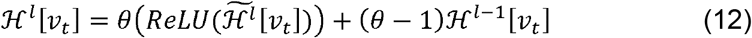

After stacking information via all *L* layers, the final node embeddings are used for downstream sample classification.

##### 3.3 Designing the loss function

The model is to solve a sample classification problem, and each sample has a known cancer label. Data imbalance is a common occurrence, which makes it challenging to accurately predict labels. To address this issue, we use Focal Loss to quantify the differences between the predicted labels and true labels (Formula (13)).

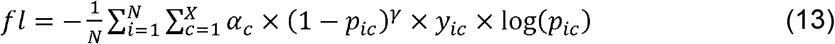

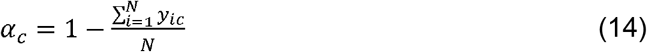

Where *N* is the total number of samples; *X* is the number of sample cancer types; *α*_*c*_ represents the weight of the *c*^*th*^ cancer type, used to balance the sample number of each class; *p*_*ic*_ is the probability that the *i*^*th*^ sample assigned to the *c*^*th*^ cancer type; *γ* is a tunable parameter (*γ* ≥ 0), and a higher value makes the model focus more on hard-to-classified samples; *y*_*ic*_ ∈ {0, 1}, and if the *i*^*th*^ sample assigned to the *c*^*th*^ cancer type, *y*_*ic*_ = 1; otherwise *y*_*ic*_ = 0. Intuitively, this function assigns more weights to nhard or easily misclassified samples than well-classified samples.

To mitigate overfitting while training the model and make the model capture more comprehensive distinguishable characteristics of each cancer type, we add a regularizer such that the final embeddings of all nodes can reflect the initial feature as much as possible, where *KL*(*,*) represents the Kullback-Leibler divergence; *A*_*M*×*N*_ is the preprocessed abundance matrix; *S*_*encoder*_ and *P*_*encoder*_ represent the final embedding matrices of species and samples, respectively.

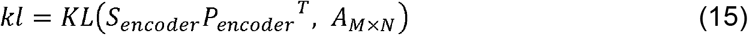

The overall loss function, denoted as *loss*, is obtained via integrating the classification loss term and the regularization term, in which *α*is the regularization factor and the default value is 0.003 considering the balance of scale between the classification loss term and the regularization term.

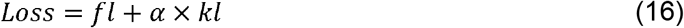

#### Step 4: Output the cancer-associated microbial communities

In the HGT model, the attention score is considered as a form of quantitative value to assess the importance of a source node to a target node. We extract the multi-head attention of each species node 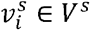 to each sample node 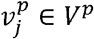, denoted as 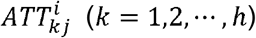, from the well-trained model, and then identify species that highly contribute to each sample based on the attention value. Given that each attention head captures distinct relations on the heterogeneous graph in their individual channel, we think the *i*^*th*^ species is of high-contribution species to the *j*^*th*^ sample as long as the attention of any of the heads 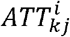 is higher than the threshold *thr*_*kj*_.

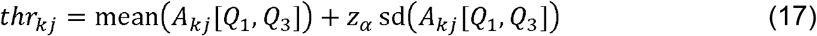

Where *A*_*k,j*_ [*Q*_1_,*Q*_3_] represents the array of all non-zero attention values between the quartile and the third quartile in the *k*^*th*^ head attention of all species to the *j*^*th*^ sample, and *z*_*α*_ is the confidence level. Here we do not use all attention in determining the threshold *thr*_*kj*_ to mitigate the impact of outliers in attention values. In this way, we can get the species that have a high contribution to each sample.

Next, the microbial community associated with a certain cancer type is inferred by detecting species with consistently high contributions to samples with the cancer type. Specifically, when identifying the microbial community associated with the *c*^*th*^ cancer type, we first collected the species with high contribution to any sample with the *c*^*th*^cancer type. Then, for each species, we count the number of samples with the *c*^*th*^ cancer type that each species significantly contributed, denoted as 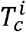 for the *i*^*th*^ species. Based on this, we can assign a *p*-value for each species (denoted as 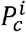 for the *i*^*th*^ species) to determine whether the species is associated with the *c*^*th*^ cancer type more often than expected purely by chance.

Our aim is to identify the microbial community associated with a certain cancer type by detecting species with consistently high contributions to samples with the cancer type. Based on the step introduced in the main text, we have obtained a species set for each sample in which each species highly contributes to the sample. In this section, we illustrated how to determine species with consistently high contributions to samples with a cancer type using the *c*^*th*^ cancer type as an example.

Suppose there are *U*_*c*_ samples with the *c*^*th*^ cancer type, denoted as 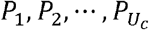 respectively. For each sample, we have obtained a set of species with high contributions, denoted as 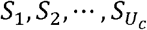. Intuitively, the microbial species associated with the *c*^*th*^ cancer type would be 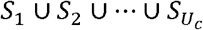. However, there may be random factors in the selection of the species associated with a sample, which will greatly increase the false positives. To mitigate this effect, we use a statistical method to select statistically significant species for cancer-associated microbial communities. Specifically, we consider the problem into the classic ball-drawing problem: *i*) Choose a number randomly from the set 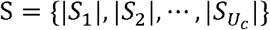, denoted as 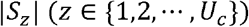. Then, remove it from *S. ii*) Draw |*S*_*z*_| balls at once from a bag with *M* different balls. Record the |*S*_*z*_| balls and then put them back. We repeat these two processes until *S* = ∅. It is important to note that we determine the number of balls selected in each experiment successively, but the order of |*S*_*z*_| should have no effect. That is to say, drawing |*S*_*i*_| or |S_*j*_| balls first are equivalent (1≤*i* ≠*J* ≤*Uc*). In this case, the probability of drawing a certain number of balls *t* or more times can be calculated, corresponding to a species highly contributing to *t* or more samples with the *c*^*th*^ cancer type. If the cumulative probability is less than 0.05, we consider the species to be significantly associated with the cancer type.

However, since the *U*_*c*_ experiments are not simply repeated, it is extremely computationally intensive to calculate the probability. To address this, we use the method of enlarging and reducing. We define 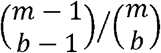 to calculate the probability of a ball being selected while randomly drawing *b* balls in a bag with *m* balls. As *b* increases, the value of 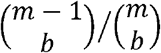 increases. We define 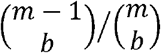 to calculate the probability of a ball not being selected. As *b* increases, the value of 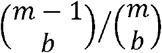 decreases. Therefore, the probability of the event “the number of a ball is drawn equal to or more than *t* times” is measured as Formula (3). This enlarging and reducing method enables us to calculate the probabilities more efficiently than directly computing them for the *U*_*c*_ experiments, allowing us to identify species significantly associated with the cancer type without incurring prohibitively high computational costs.

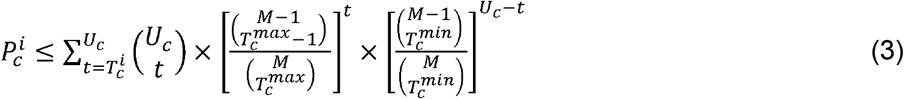

Where *U*_*c*_ is the total number of samples with the *c*^*th*^ cancer type. *M* is the number of all species. 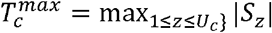 and 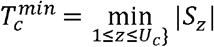 represent the maximum and minimum number of samples with the *c*^*th*^ cancer type that each species significantly contributed to, respectively. If the value on the right-hand side of the Formula (3) is less than 0.05, the species will be regarded as significantly contributing to the *c*^*th*^ cancer type.

### The evaluation of cancer classification performance

Several machine learning methods, such as Lasso, Enet, RF, SVM, and MIIDL, were used to compare classification performance. Lasso, Enet, RF, and SVM were implemented by the MetAML software^23^. MIIDL was implemented via the python package miidl 0.0.5. All of them were run on the Pitzer cluster of the Ohio Supercomputer Center with memory usage set to 300GB. We optimized the tunable parameters of each method using a grid search. We tuned multiple parameters for the machine learning methods.

For MIIDL, we tried different parameter sets, including: (1) qc: 0.0001, 0.001, 0.1, and 0.3; (2) Normalization: log_2_(*x* + 1), log2(*x*), ln *x*, ln(*x +* 1), mean, median, z-score, and none; (3) Imputation: knn, mean, median, minimum, and none; (4) Epoch: 100, 200, 400, and 600. Finally, we selected the parameter combination that performed the best on the TCMA dataset, which has qc: 0.00001; normalization: loge; imputation: mean; and epoch: 600.

For the four methods (Lasso, Enet, RF, and SVM), we used the MetaML software and optimized the classifiers using GridSearchCV for the best model. The parameter set included: (1) i: lasso, enet; (2) s: f1_macro; roc_auc; (3) r: 10, 20, 40. Other parameters, such as the number of features, were used in the default ranges. Finally, the parameters used for the four methods were as follows: Lasso: i enet s f1_macro r 10; Enet: i enet s f1_macro r 10; RF: i enet s f1_macro r 40; SVM: i enet s f1_macro r 10.

Accuracy, precision, recall, and F1-score were used to evaluate the classification performance, in which accuracy was calculated based on the overall prediction results, and the others were calculated using the average across five cancer types.

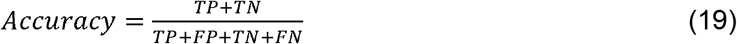

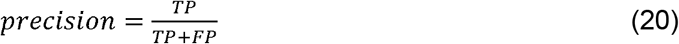

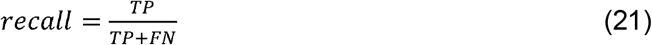

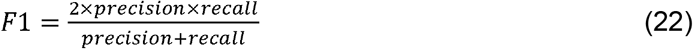

Where the true positive (TP) represents the number of samples with the cancer that are correctly classified; the false positive (FP) represents the number of samples that are incorrectly classified with the cancer; the true negative (TN) represents the number of samples without the cancer that are correctly classified; and the false negative (FN) represents the number of samples with the cancer that are incorrectly classified.

### The evaluation of reproducibility performance

To assess the reproducibility of identified cancer-associated microbial characteristics, we stratified randomly subsampling a certain proportion (denoted as *r*) of samples from the original dataset to form a ‘new’ dataset. The random selection repeated *K* times and generated *K* datasets, denoted as 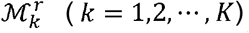. Then, the method for cancer-associated microbial characteristics identification was applied to 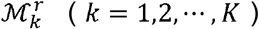, respectively. The identified species of the *c*^*th*^ cancer type was represented as 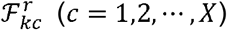. Then, for each cancer type, we used the Jaccard index to assess the similarity between the characteristics identified by each of the two datasets. The average 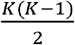 pairwise similarities,*RJ*_*c*_, was used as the reproducibility index for the *c*^*th*^ cancer type. The average of the reproducibility indexes across all cancer types, *RJ*, was used to assess the overall reproducibility of the identified characteristics, and a higher value corresponds to a higher reproducibility.

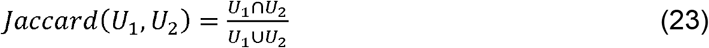

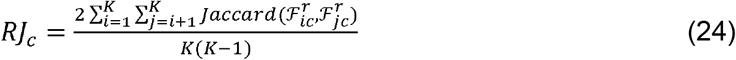

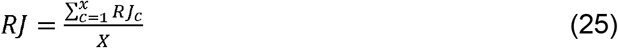

Here, we set *K* = 50and *r* ranging from 80% to 95% with a step of 5% to assess the reproducibility. The competing methods include a statistics-based method, LEfSe, and a network-based method, NetMoss, as well as the five machine learning methods (Lasso, Enet, RF, SVM, and MIIDL). LEfSe was implemented using the published source code; NetMoss was implemented using the R package NetMoss 0.1.0.

## RESULTS

### MICAH consistently outperformed other methods in terms of classification performance

We first assessed the performance of MICAH in differentiating between different cancer types using intratumoral microbial species abundance data from TCMA^10^. We expected the proposed framework can discriminate well among the five cancer types, suggesting the capability of capturing microbial features associated with different cancer types. Ten-fold cross-validation was used here, and results showed that the accuracy, precision, recall, and F1-score of MICAH across the five cancer types were 0.8053, 0.7539, 0.7304, and 0.7318, respectively (**Fig. 2a**). Observation on individual cancer types found the precision, recall, and F1-score varied widely across different cancer types (**Table 1**). The results of ESCA and READ were significantly lower than others. This may be caused by the variation in sample size since the classification performance of each cancer type suggested a clear positive correlation with the sample size. We compared MICAH with several commonly used methods for identifying disease-associated microbiome by solving classification problems, including traditional machine learning methods (Least absolute shrinkage and selection operator, abbr. Lasso; Elastic net, abbr. Enet; Random forest, abbr. RF; and Support vector machine, abbr. SVM) and a GNN method, MIIDL. The four metrics, i.e., accuracy, precision, recall, and F1 score, were calculated. As MIIDL is a python package without publicly available source code, it only outputs classification accuracy, so we only show this metric for it. Results suggested MICAH has the best performance in cancer type classification, followed by MIIDL, RF, Lasso, Enet, and SVM (**Fig. 2a**). We acknowledge there were many other algorithms for cancer classification^35-37^. But they focus more on classification rather than downstream cancer-associated microbiome analysis and lack interpretation ability^37^. We thus did not include them in this benchmarking comparison.

**Table 1.** The classification performance of the five cancer type.

|  | COAD | ESCA | HNSC | READ | STAD |
| --- | --- | --- | --- | --- | --- |
| **Precision** | 0.8093 | 0.4339 | 0.8471 | 0.6115 | 0.7902 |
| **Recall** | 0.7987 | 0.45 | 0.767 | 0.77 | 0.741 |
| **F1-score** | 0.7962 | 0.4224 | 0.8008 | 0.667 | 0.7542 |

**Fig. 2.**
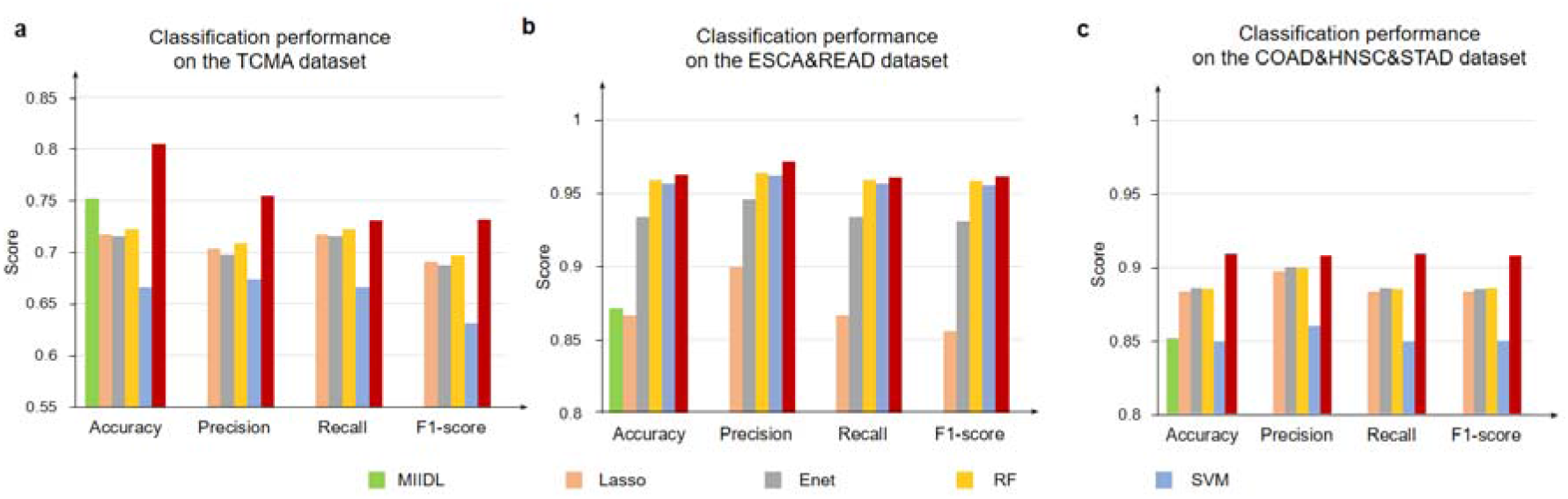
Results of classification performance. a. Accuracy, precision, recall, and F1 score of MICAH, Lasso, Enet, RF, and SVM. b & c. Classification performance of two batches (the ESCA&READ dataset and the COAD&HNSC&STAD dataset).

To assess the generalizability of MICAH across datasets, we split the five cancer types into two batches based on a previous study^3^. This was implemented based on sample size, generating two new datasets: one containing ESCA and READ samples (called the ESCA&READ datasets), and the other containing COAD, HNSC, and STAD samples (called the COAD&HNSC&STAD dataset). This is also conducive to exploring the classification performance of different methods in the case of relatively balanced data. We applied MICAH and the five methods above on the two datasets and found that the classification performance of all these methods improved. Not surprisingly, MICAH showed consistently strong performance in distinguishing cancer types based on intratumoral microbial characteristics (**Fig. 2b & c**), suggesting its high generalizability.

### MICAH showed superior reproducibility than other methods in identifying cancer-associated microbial communities

We further measured the reproducibility of identified communities from MICAH. Ideally, the true characteristics of each cancer type should not be sensitive to sample variations. Data perturbation, such as changing or taking out a small number of samples, should not significantly alter the identified microbial characteristics. To test this, we randomly selected a certain proportion of samples from the original TCMA dataset, forming a ‘new’ dataset. This process was repeated 50 times and generated 50 datasets with a small fraction of differential samples. The intratumoral microbial characteristics were identified by MICAH, as well as seven other competing methods, i.e., Lasso, Enet, RF, SVM, MIIDL, LEfSe (a representative statistics-based method), and NetMoss (a representative network-based method). A reproducibility index was then used to measure the similarity of identified characteristics across the 50 datasets (METHODS). Results indicate that MICAH has higher reproducibility than other methods for small sample variations (**Fig. 3a**). Nevertheless, when the proportion of subsampling was less than 85%, the reproducibility of MICAH is slightly lower than the statistics-based method, LEfSe. This may be due to the deep-learning model’s inherent limitations in sample size requirements. Further observation on the first 20 identified species found the concentration of the plots for the ranks of identified microbes from MICAH was highest around the 45° line (R^2^ = 0.8216), followed by LEfSe (R^2^ = 0.7289) and RF (R^2^ = 0.6689), while those of other methods seem spread with R2 < 0.50 (**Fig. 3b**). This indicated that MICAH retained the ranks of the selected characteristics well. Our findings demonstrate that MICAH is highly reproducible in identifying cancer-associated microbial characteristics.

**Fig. 3.**
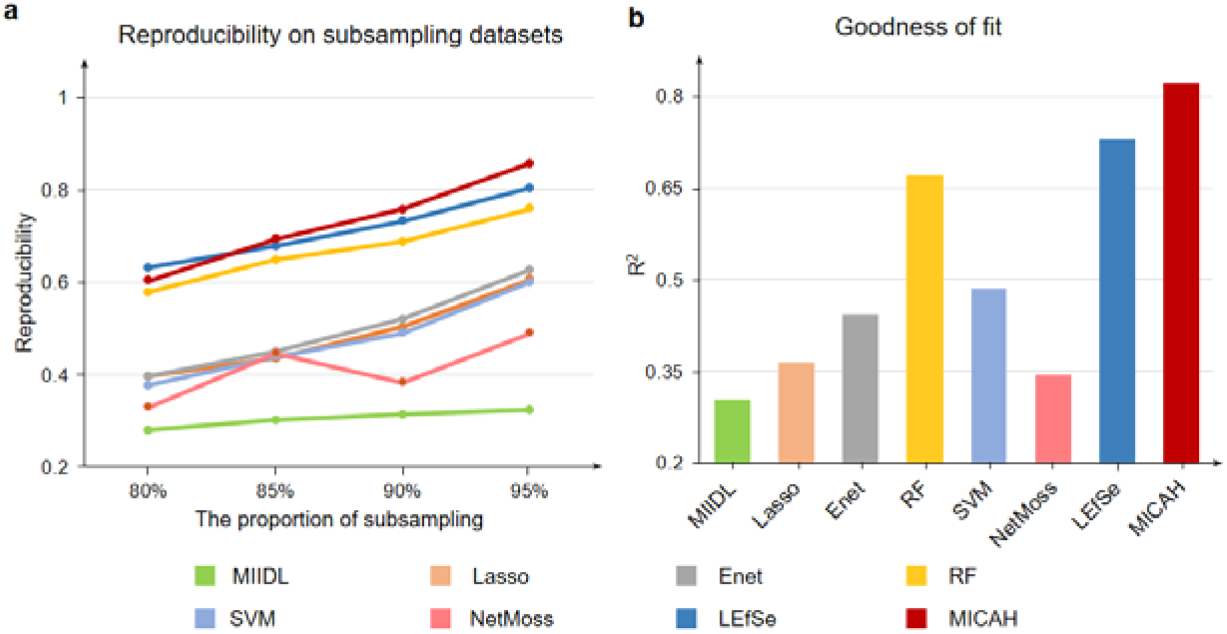
Results of reproducibility performance. **a**. Reproducibility of MICAH and seven other competing methods in different subsampling datasets of TCMA. **b**. Goodness of fit with the line y=x of rank plots for the eight methods.

## DISCUSSION

Although the cancer-associated microbiome has been extensively studied for decades, the relationships between intratumoral microbes and cancer are far from fully understood^38^. To this end, we developed the MICAH framework to identify cancer-associated intratumoral microbial communities. MICAH has three key features: (*i*) an explainable deep learning framework to infer cancer-associated intratumoral microbial communities; (*ii*) abstracting metabolic and phylogenetic relationships among all species in a graph representation learning framework to systematically reflect the intricate interactions within microbial communities; and (*iii*) introducing the attention mechanism for holistic interpretation of the relevance between microbial species and sample phenotypes. Using MICAH, we have successfully identified intratumoral microbial characteristics of five cancer types. The success partially comes from the power of GNN for the modeling of complex relationships among samples, microbes, and cancer types, including their local and global dependencies. GNN is robust to noisy or incomplete data due to their ability to incorporate information from neighboring nodes in the graph. In addition, the heterogeneous graph transformer framework of GNN is computationally efficient and scalable to large graphs. It took about an hour to get final results on the TCMA dataset. We believe that MICAH can contribute to the understanding of complex microbial communities and guide biologists in understanding cancer pathogenesis. Furthermore, our framework can be extended to identify microbial signatures associated with different diseases and environmental exposure.

However, this study has some limitations. First, MICAH relies on an existing database to assess metabolic relationships between species, which may not fully reflect true metabolic activities within the community in specific cancer. Admittedly, it is a way to assess metabolic relationships to compensate for the lack of metabolic data. This issue can be addressed if the user can provide metabolic data for species-species relationships assessment. Furthermore, the metabolic and phylogenetic relationships among species in this study are only a simplified representation, and their directionality and strength have not been considered. Moving forward, improving the representation of these relationships will be a key focus in achieving a more comprehensive understanding of how species interact and influence each other within microbial communities. Second, mechanistic interpretation of identified microbial communities to cancer remains difficult based only on the microbial data from bulk tissue sequencing. Spatial sequencing and single-cell sequencing data provide a more comprehensive understanding of the spatial distribution of microbes within human tissue and their interactions with specific cell types^39,40^. By leveraging these data, researchers can gain insights into the microbial community structure, functions, and their relationships with host cells, leading to a deeper understanding of the role of intratumoral microbiome in cancer development and progression. Third, we focus on cancer-associated microbial species communities, while it is insufficient to understand microbial communities in species-level resolution^41^. Considerable genomic and phenotypic variabilities have been observed within species^41^, and several species have both pathogenic and commensal strains^42^. Therefore, a more detailed characterization of microbial communities for within-species variation is necessary for cancer-associated microbial signature exploration. Additionally, time-series data should be introduced in future studies to carry out in-depth longitudinal analysis of dynamic microbial communities and their interactions with cancer. A relative temporal encoding strategy can be used to model temporal dependencies in a GNN model, thereby capturing longitudinal dysbioses in cancer and improving the understanding of cancer development.

## DATA ACCESS

The dataset supporting the conclusions of this article is available in the TCMA database (https://doi.org/10.7924/r4rn36833). The source code of MICAH is freely available at https://github.com/OSU-BMBL/micah.

## AUTHOR CONTRIBUTIONS

Q.M. and B.L. conceived the basic idea. Z.L., Y.S., and X.W. carried out the computational analysis and data interpretation. L.Z. and A.M. designed and drew the figures. D.S., Q.M., D.X., and B.L. polished the manuscript. All authors read and approved the final manuscript.

